# Self-supervised learning of probabilistic prediction through synaptic plasticity in apical dendrites: A normative model

**DOI:** 10.1101/2021.03.04.433822

**Authors:** Arjun Rao, Robert Legenstein, Anand Subramoney, Wolfgang Maass

## Abstract

Sensory information is processed by the brain not in a simple feedforward fashion. Rather, bottom-up inputs are combined in pyramidal cells of sensory cortices with top-down information from higher brain areas that arrives through synapses in apical dendrites. The exact functional role of these top-down inputs has remained unknown. A promising abstract model posits that they provide probabilistic priors for bottom-up sensory inputs. We show that this hypothesis is consistent with a large number of experimental about synaptic plasticity in apical dendrites, in particular with the prominent role of NMDA-spikes. We identify conditions under which this synaptic plasticity could approximate the gold standard for self-supervised learning of probabilistic priors: logistic regression. Furthermore, this perspective suggests an additional functional role for the complex structure of the dendritic arborization plays: It enables the neuron to learn substantially more complex landscapes of probabilistic priors.

## 1 Introduction

A key open problem in neuroscience and cognitive science is understanding how bottom-up sensory information and internally generated top-down information from higher brain areas are integrated in the computations of laminar cortical microcircuits [Vezoli et al., 2021, Larkum, 2013]. Several abstract theoretical frameworks have been proposed for understanding the organization of these computations, for example that top-down information from higher areas produces probabilistic (Bayesian) priors for bottom-up information [Lee and Mumford, 2003], or internal reconstructions (predictions) of sensory inputs [Rao and Ballard, 1999]. But it remains difficult to relate these abstract theories to experimentally observed processes in cortical microcircuits. Experimental data clarify where topdown and bottom-up information meet in generic cortical microcircuits: in pyramidal cells on layers 2/3 and 5. More precisely, top-down inputs reach these cells primarily through synapses on their apical dendrites in superficial layers, especially in layer 1 [Larkum, 2022], whereas bottom-up information primarily targets basal dendrites. Hence we need to understand the plasticity processes which shape the impact and information content of this top-down synaptic input. It has been proposed by [Shin et al., 2021] that two sequential phases of learning should be distinguished: During the first phase, which takes substantial time and primarly involves synaptic plasticity in basal dendrites, these pyramidal cells are grounded in the most prevalent statistical features of the environment. During the second phase the activity of these neurons is linked to internal models, using the putative predictive role of apical dendrites. Experimental data show in fact that plasticity rules for synapses in apical dendrites differ from the more commonly considered Hebbian and STDP-like rules for basal dendrites. In particular, local NMDA spikes in the apical tuft of pyramidal cells play an essential role for this plasticity, see e.g. [Augusto and Gambino, 2019, Stuyt et al., 2021, Larkum, 2022] for recent reviews. Related to the important role that NMDA spikes, which typically last 100ms and longer, play for synaptic plasticity in apical dendrites, unlike STDP for basal dendrites the precise temporal relation between these NMDA spikes and other salient events, appears to be less relevant for inducing synaptic plasticity. But also the functional goal of these plasticity processes in apical dendrites remains unclear. Hence we are addressing here the question to what extent these processes, and in particular dendritic spikes, could support self-supervised learning of a prediction of bottom-up inputs to the neuron.

Self-supervised learning of probabilistic predictions requires rules for synaptic plasticity that differ from the more commonly considered rules for classification learning: Predictions are inherently uncertain, i.e., they turn out to be sometimes correct and sometimes not. Most rules for synaptic plasticity that are suitable for classification learning, such as the perceptron learning rule [Moldwin and Segev, 2020], do not perform well if the labels of training examples are inconsistent -which is the generic scenario for learning probabilistic predictions of uncertain binary events. Furthermore, for learning probabilistic predictions, a plasticity rule has to converge to a probability, rather than to a guessed classification. A gold standard for such learning is logistic regression [Bishop, 2006], since this method learns a probabilistic prediction that is, under some constraint on the mathematical form of the prediction, theoretically optimal. We examine to what extent rules for synaptic plasticity in apical dendrites that are consistent with experimental data can approximate logistic regression. More precisely, we derive an arguably optimal approximation to logistic regression through synaptic plasticity in apical dendrites that we term Dendritic Logistic Regression (DLR). Numerous experimental data on synaptic plasticity in apical dendrites are consistent with DLR. In particular, dendritic spikes (NMDA spikes) play a key role for DLR.

Our theoretical framework holds for any number of top-down inputs, however in two of our demonstrations (in Fig. 2, 4), we consider just the case of 2-dimensional top-down inputs **x**, because learnt landscapes of probabilistic priors in the space of top-down inputs can easily be visualized in this case. We first present the theoretical framework for the case of a single dendritic branch in the apical tuft of a pyramidal cell. The extension to the case of several dendritic branches is presented subsequently. Since experimental data point to two types of interactions between dendritic branches in the apical tuft of different types of neurons, a saturating summation [Naud et al., 2014, Poirazi et al., 2003, Gooch et al., 2022], as well as a soft XOR for some types of pyramidal cells on layer 2/3 of the human cortex [Gidon et al., 2020], as predicted by the model of [Gorski et al., 2017], we analyze functional consequences of DLR for both cases. It turns out that both types of neuron models support DLR, but facilitate learning of different types of probability landscapes. Since either type of probability landscape is useful in some scenarios and less in others, the resulting learning theory for apical dendrites supports the co-existence of different subtypes of pyramidal cells in layer 2/3 [Planert et al., 2021] with different interactions between apical dendrites, since this guarantees that good probabilistic predictions can be learnt in each scenario by at least some of the neurons.

**Figure 1:**
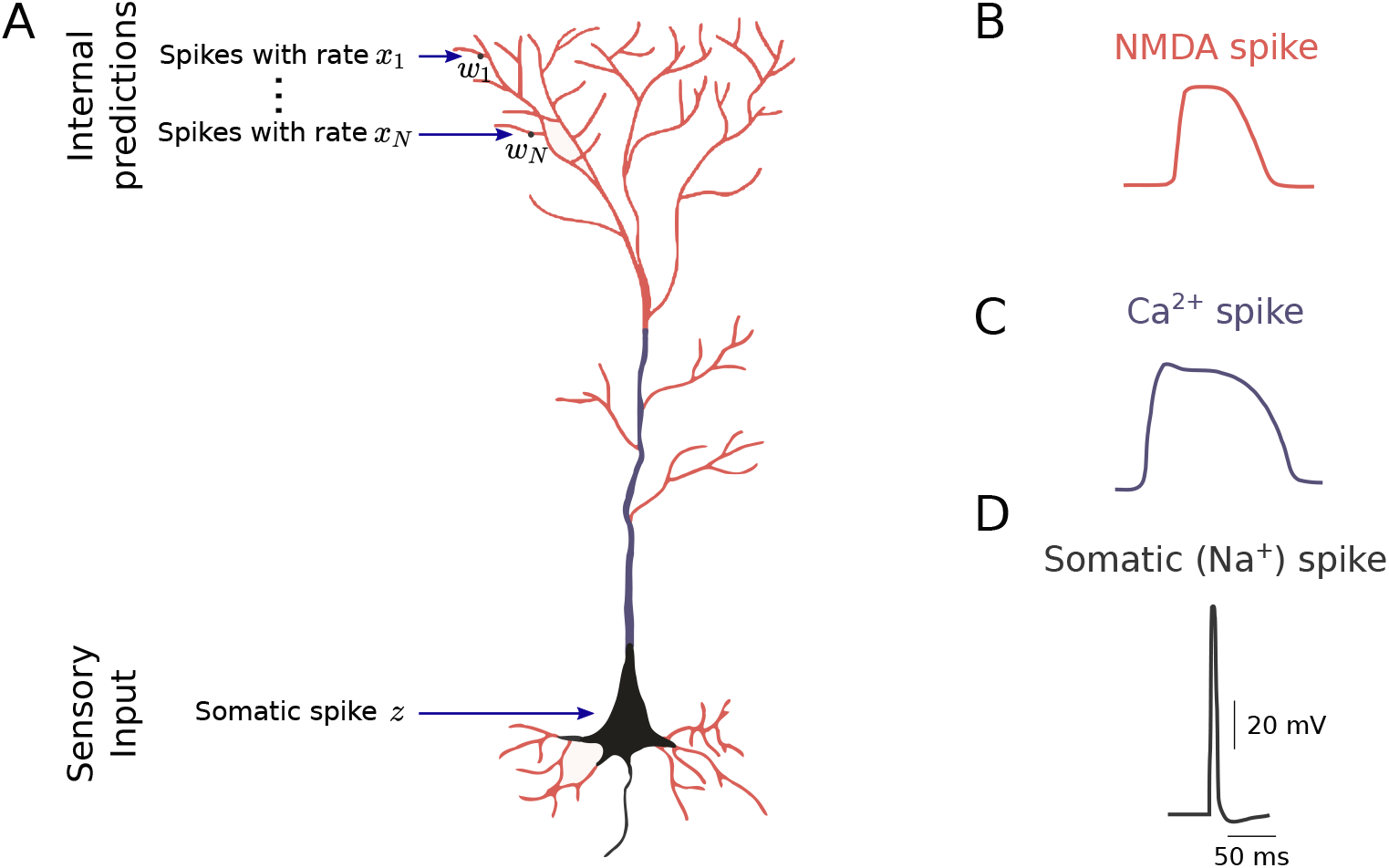
Scheme of a pyramidal cell with apical dendrites, dendritic spikes, and somatic spikes. **A)** Generic model of a thick tufted pyramidal cell in layer 2/3 or layer 5 of a generic laminar cortical microcircuit. It integrates two different inputs streams, a top-down stream that arrives in its apical dendrites (shown at the top), and a bottom-up stream that arrives in basal dendrites and influences the generation of somatic spikes. see [Larkum, 2013, Augusto and Gambino, 2019, Larkum et al., 2022]. Synaptic inputs from neurons in higher brain areas with firing rates *x*_1_, …, *x_N_* are integrated with synaptic weights *w*_1_, …, *w_N_* in the apical tuft. We propose a normative learning model for synaptic plasticity of these weights that aims at learning the probability of a co-occurring somatic spike. **B)** NMDA spikes in apical dendrites play a major rule in this normative model. **C)** In view of the larger distance between apical dendrites and the soma for a oyramidal cell on layer 5, information about a somatic spikes reaches apical dendrites through an intermediate signal, a Ca^2+^-spikes, which is generated in the dendritic trunk (blue region in panel A). **D)** A somatic spike. This figure is adapted from [Augusto and Gambino, 2019].

**Figure 2:**
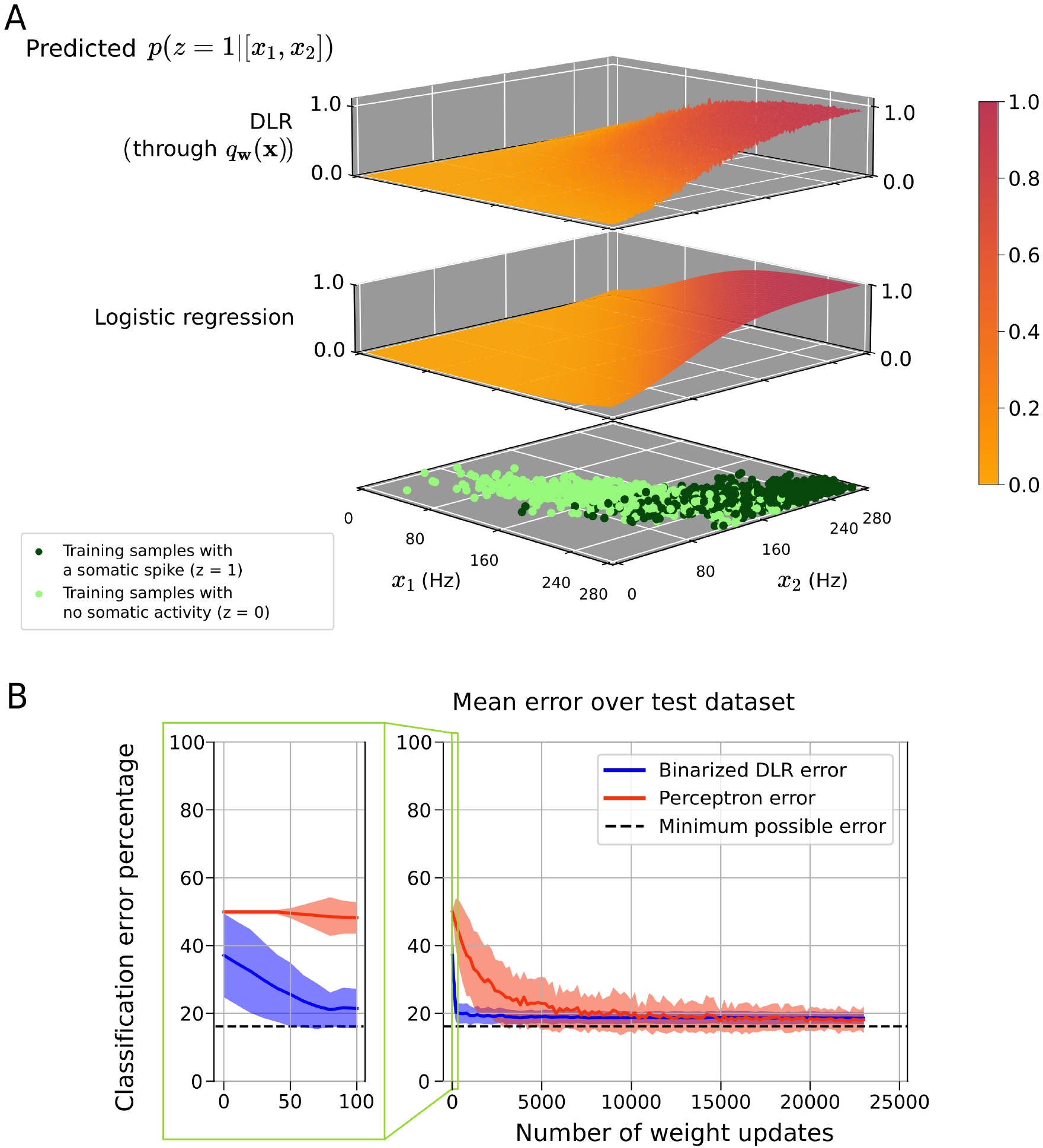
Illustration of the challenge to learn probabilistic predictions, and performance of the gold standard logistic regression as well as a biologically plausible approximation (Dendritic Logistic Regression: DLR) on a simple benchmark task. **A)** Bottom row: Top-down inputs **x** are generated by two overlapping Gaussians. The **x** from one of the Gaussians are accompanied by somatic spikes (dark green points denote some samples), the other Gaussian generates **x** that are not accompanied by somatic spikes (like green sample points). Middle row: Probability that a top-down input **x** is accompanied by a somatic spike, estimated through logistic regression. Top row: DLR produces almost equally good estimates for the same task (mean difference 0.045). **B)** The perceptron learning rule, as considered in [Moldwin and Segev, 2020], can only learn binary classifications, no probabilities. Hence the performance of the preceptron learning rule (red curve) and DLR (blue curve) can only be compared if one ignores the additional information that DLR learning provides through in the form of a probability, and rounds its learnt probabilities to 0 or 1 (Binarized DLR). An interesting difference between these two approaches for learning classifications is that DLR converges in much fewer weight updates to provide close-to-optimal classifications. Mean and variance of the error are shown for 100 learning episodes with independently drawn initial weights and data points.

For simplicity, we will refer to the dendritic branches in the apical tuft of pyramidal cells in layer 2/3 or 5 just as apical dendrites.

## 2 Results

### 2.1 A normative model for self-supervised learning of probabilistic predictions through synaptic plasticity in an apical dendrite

The main principle of the common hierarchical Bayesian inference model [Lee and Mumford, 2003] for integrating top-down information from higher brain areas into processing of bottom-up input streams in the cortex is the following: The top-down information **x** is used on each level of the cortical hierarchy to produce a prior *p*(*z*|**x**) for the bottom-up input to that level. Anatomical data show that these two information streams meet in pyramidal cells on layers 2/3 and 5 of the generic cortical microcircuit. More precisely, the top-down information arrives primarily on apical dendrites and bottom-up information primarily on basal dendrites of pyramidal cells. For an individual pyramidal cell, the Bayesian inference model of [Lee and Mumford, 2003] amounts to the challenge of estimating the probability that input from basal dendrites causes a somatic spike, given the firing rates **x** of neurons in higher cortical areas that form synapses on its apical dendrites. We denote the presence or absence of a somatic spike (within some suitable time-window) by a binary variable *z*. Obviously, the synaptic weights **w** = [*w*_1_, *w*_2_, …, *w_N_*] in apical dendrites are essential for transforming firing activity in higher brain areas into a prediction of somatic spikes of an individual neuron. Logistic regression [Bishop, 2006], see Methods, provides a gold standard for this probabilistic prediction, since it is theoretically optimal among all predictions that apply the sigmoid activatation function in order to map the weighted sum 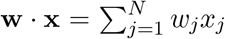 into the range of probabilities, i.e., to values between 0 and 1.. We examine to what extent this gold standard can be approximated by synaptic plasticity rules for apical dendrites.

Experimental data show that local NMDA spikes are essential for synaptic plasticity in apical dendrites [Gambino et al., 2014]. Full knowledge of the underlying molecular processes is not yet available. But it has been shown in [Stein et al., 2021] that non-ionotropic NMDA receptor signaling plays a key role. Furthermore, it was shown there that the presence or absence of coincident Ca^2+^-influx decides between LTP and LTD. We use the binary variable *y* to denote the presence or absence of an NMDA spike in the apical dendrite. We denote the probability of an NMDA spike in an apical dendrite, in dependence on the synaptic input **x** that it receives, by *q*_**w**_(**x**). We examine whether this term can be tuned through biologically plausible synaptic plasticity rules to provide an estimate of the probability *p*_target_(*z* = 1|**x**) of a somatic spike of the neuron. We use for the term *q*_**w**_ (**x**) the simple formula

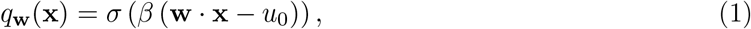

where 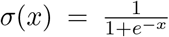 is the standard sigmoid function, and *β* is a constant scaling factor. The threshold *u*_0_ can be interpreted in biophysical terms as the minimal membrane voltage that is required to remove the *Mg*^2+^ block of NMDA receptors. The term *q*_**w**_(**x**) can then be related to the dendritic excitability of an apical dendrite, more precisely to the probability of generating an NMDA spike in response to top-down input **x**. According to the experimental data reviewed in [Stuyt et al., 2021, Gonzalez et al., 2022] this dendritic excitability changes during learning, and is an important factor in the induction of LTP. Furthermore, the resulting modified dendritic excitability plays an important computational role: it tunes the perceptual threshold [Takahashi et al., 2016, Takahashi et al., 2020].

In order to derive a plasticity rule for apical dendrites from the goal of predicting *p*_target_(*z*|**x**) through the probability of an NMDA spike in the dendrite, we use the common cross entropy loss as a measure for the distance between the ground truth probability and the estimate *q*_**w**_(**x**). This loss function is convex, and hence can be optimized through stochastic gradient descent. This stochastic gradient descent yields the following rule for synaptic plasticity for the weights **w** of synapses onto the apical dendrite:

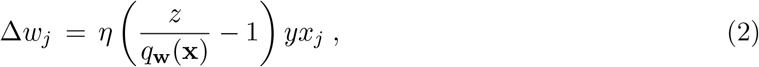

where Δ*w_j_* is the change of the weight of synapse *j* and *η* > 0 is a constant scaling factor. The rule can equivalently be described in the form of Table 1. We show in Section 3.2, that this weight update performs stochastic gradient descent for the previously described cross entropy loss. Since this stochastic gradient descent is guaranteed to converge to the minimal value of this loss function. Furthermore, since this minimal value is the same as the value that can be achieved through theoretically optimal logistic regression for approximating the target value *p*_target_(*z*|**x**), it gives rise to a normative model. We refer to this normative plasticity rule as Dendritic Logistic Regression (DLR).

**Table 1:** Updates for synaptic weight *w_j_* according to the Dendritic Logistic Regression (DLR) rule: The firing rates of presynaptic neurons in higher brain areas are denoted by **x**, and *z* represents a somatic spike (or a global plateau potential in some types of neurons).

| NMDA spike $y$ | Somatic spike $z$ | weight update $\Delta \mathbf{w}$ |
| --- | --- | --- |
| 0 | 0 | 0 |
| 0 | 1 | 0 |
| 1 | 0 | $-x_j$ |
| 1 | 1 | $\frac{x_j}{q_{\mathbf{w}}(\mathbf{x})} - x_j$ |

DLR is consistent with the experimental data of [Gambino et al., 2014, Stein et al., 2021] insofar as synaptic plasticity only occurs in the presence of an NMDA spike, i.e., *y* = 1. Furthermore, consistent with the results of [Stein et al., 2021], the decision between LTP and LTD depends on the presence of a signal that causes Ca^2+^-influx in temporal proximity, which can be caused in L2/3 pyramids by a backpropagating action potential, and in L5 pyramids by a Ca^2+^-spike that is triggered when significant depolarization in the apical tuft coincides with a somatic spike [Larkum, 2013, Larkum et al., 2022]. In addition, DLR predicts that the amplitude of LTP is scaled by the term *q*_**w**_(**x**) = *σ* (*β* (**w** · **x** – *u*_0_)), i.e., by the current excitability of the dendrite for NMDA spikes, reducing the amplitude of LTP if this excitability is already fairly high.

#### 2.1.1 A simple illustration of the capability of DLR to learn probabilistic predictions

We consider two clusters of firing patterns **x** of neurons in higher brain areas that project to an apical dendrite of a pyramidal cell. One of them models activity patterns **x** that were accompanied by a somatic spike (dark green points at the bottom of in the bottom row of panel A in Fig. 2), and the other models activity patterns **x** that were not accompanied by a somatic spike (light green points in the bottom row of panel A in Fig. 2). Importantly, in order to model the inherent uncertainty of predictions we consider the case that these two clusters of firing patterns **x** of neurons in higher brain areas overlap, i.e., the same **x** is sometimes accompanied by a somatic spike and sometimes not. More precisely, each cluster is generated by a Gaussian 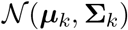, with parameters (***μ***_0_, **Σ**_0_) and (***μ***_1_, **Σ**_1_), and these two Gaussians overlap. In order to facilitate visualization, we consider here just two-dimensional firing patterns **x**, but the same principle applies for inputs of arbitrary dimension from higher brain areas.

Fig. 2A shows that the apical dendrites learn through DLR to predict via *q*_**w**_(**x**) quite precisely the probability that an input pattern **x** is accompanied by a somatic spike. In fact, one sees that the resulting prediction performance was very similar to that of theoretically optimal logistic regression. The probabilistic prediction capability that results from DLR can be measured through the negative log likelihood (NLL) in nats (lower is better) for the training data. Across a 100 independent training runs, it achieved an average of 0.384 nats, compared to 0.358 nats achieved through logistic regression. We show in Fig. S1 of the Supplement that DLR also learns close to optimal prediction probabilities if these approach nowhere the extreme values of 0 or 1.

A direct comparison of convergence properties of the perceptron learning rule and DLR requires us to binarize the probabilistic prediction that is learnt through DLR, since the perceptron learning rule can only learn to produce binary classifications. However, the two point clouds are not linearly separable due to the overlap of the Gaussians which generate them, and hence the perceptron learning rule (as investigated in [Moldwin and Segev, 2020] struggles with separating them, due to the inconsistencies of classifications in the training set, as indicated by the red trace in Fig. 2B. In contrast, the blue trace in Fig. 2B shows that binarized DLR converges quite fast to an optimal linear separation of the two point clouds. This large performance difference between perceptron learning and DLR might seem surprising. The perceptron weight update rule (see equation 21) has dependencies only based on the binary condition of whether **w** · **x** is greater or lesser than *u*_0_. In comparison, the more fine-grained dependence of the DLR on dendritic excitability *q*_**w**_(**x**) (equation 1) greatly improves the convergence properties.

#### 2.1.2 Predicting future sensory input caused by a moving object

Our brains are very good at estimating from the current position, speed, and movement direction of a moving object the probability that this object will be at a given later time point *T* at a particular position. This is a non-trivial prediction task since estimates of initial position, movement speed and direction are inherently noisy, see the cartoon in Fig. 3B. We present a simple model which shows that these probabilistic predictions can be acquired through DLR. We assume that the receptive fields of a 2D array of neurons cover the movement field, and for simplicity, we represent the moving object by a linearly moving disk. The neurons in this array could for example be neurons in a visual or somatosensory area. We furthermore assume that these neurons receive from higher brain areas the estimated position, speed, and movement direction of a moving object at time 0, see Fig. 3A. We model the uncertainty of these estimates by adding Gaussian noise to an ideal estimate of current speed and movement direction.

**Figure 3:**
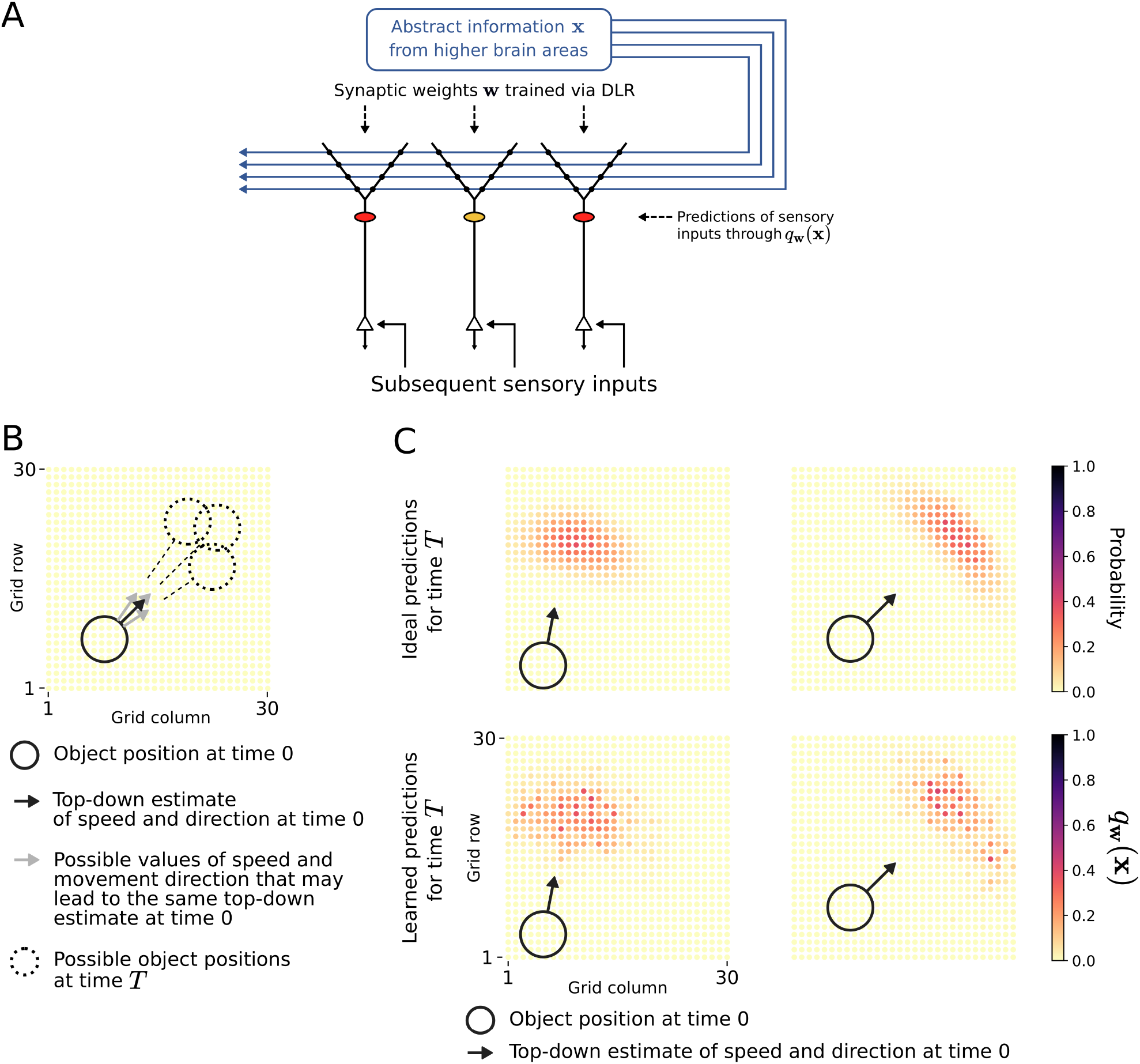
Performance of DLR for self-supervised learning to predict whether the receptive field of a neuron will be covered by a moving object. **A)** Schema of the simple network architecture, a 30 × 30 neuron sheet of neurons with apical dendrites that each have their own receptive field, but receive the same top-down input **x**. **x** denotes in this case estimates of the initial position, speed and movement direction of some object. When the object covers at a later time T, the prediction span, the receptive field of a neuron it causes a somatic spike. Each neuron learns through synaptic plasticity in its apical dendrites as estimate of the probability *q*_**w**_(**x**) that this will happen (color code for individual predictions as in panel C. **B)** Because of uncertainty in the estimates of the initial position, speed, and movement direction of the object, this top-down input can only give rise to probabilistic estimates of the position of the object at time T. **C)** For two instances of this scenario we show in the lower row for each neuron in the 30 × 30 array its individual probabilistic prediction *q*_**w**_ (**x**) that it generates in its apical dendrites, after self-supervised learning with DLR. For comparison, in the upper row, we show for each grid position, the actual probability that it will be covered at time *T* by the moving object, computed using the knowledge of the full distribution of noisy estimates for time 0 (see Section 3.6.3, Methods).

The underlying self-supervised learning task is specified as follows. Each neuron in this 2D array receives the estimates of position and velocity at time 0, encoded as spike rates **x**, that arrive at its apical dendrites from higher brain areas. Furthermore, each neuron fires a somatic spike *z* when its receptive field is covered by the moving object at time *T*. Hence we can interpret the target distribution *p*_target_(*z* = 1|**x**) as the probability that the object will be positioned over the receptive field of a neuron at time *T*, given the position and velocity at time 0.

According to the underlying theory, once the weights have been learned via *DLR*, *q*_**w**_ (**x**) approximates for each neuron in the 2D grid *p*_target_(*z* = 1|**x**). The learnt probabilities are shown in the lower row of Fig. 3C for two different movement scenarios. A comparison with the probabilistic ground truth in the upper row of Fig. 3C shows that these learnt predictions are close to optimal for this task. Note that the probabilistic ground truth in the upper row was computed on the basis of full knowledge of the distribution of the noisy estimates of speed and direction at time 0.

### 2.2 Extension to the case of synaptic plasticity in multiple apical dendrites

When there are several dendritic branches, the question arises as to how the contributions from different branches are combined for the prediction *q*_**w**_(**x**), that is in our model the probability of a dendritic spike arising from apical dendrites. Here, we denote the weights and presynaptic firing rates for synapses in each dendritic branch *B* by 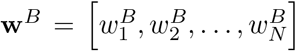 and 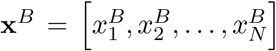 respectively, and the weights and presynaptic firing rates across all branches as **w**, **x** respectively. As in the single dendrite case, the goal of synaptic plasticity in apical dendrites is in our model that *q*_**w**_ (**x**) learns to become a good estimate of the target probability *p*_target_(*z* = 1|**x**).

In order to arrive at a model for the probability *q*_**w**_(**x**) of a dendritic spike arising from the apical dendrite, we assume that contributions *f^B^* from each branch *B* are summed and followed by a nonlinearity Ψ:

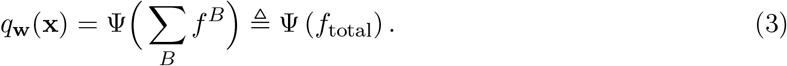

Consistent with the single-branch model (equation 1), the contribution of each branch is given by a sigmoidal function of its weighted input rate, i.e.

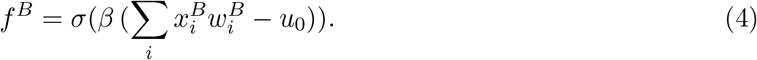

For the nonlinearity Ψ we consider two possibilities. According to the classical view [Poirazi et al., 2003, Tzilivaki et al., 2019, Jadi et al., 2014], the dendritic activity is given by a saturating nonlinearity. Hence, as a first possibility, we model Ψ as a sigmoidal function in Subsection 2.2.1 below. More recently, a more complex relationship was found for single human neurons [Gidon et al., 2020] which will be considered in Subsection 2.2.2.

#### 2.2.1 Saturating model

In the first possibility, taking into consideration the thresholding and saturation dynamics of pyramidal neurons [Naud et al., 2014, Jadi et al., 2014], we chose the sigmoidal shape for Ψ to get

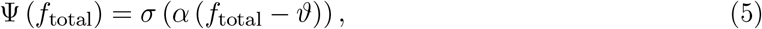

where *α* and *ϑ* are suitable constants. Note that by choice of *α* and *ϑ*, one can model different types of (soft-) logic operations. A value of *ϑ* less than 1 implements a soft OR operation (a minimum of one branch needs to be active to get a high probability output). A value of *ϑ* close to the number of branches implements a soft AND, and in-between we obtain a general threshold condition on the number of active branches. See Fig. 4B (left and middle) for an example with two branches.

**Figure 4:**
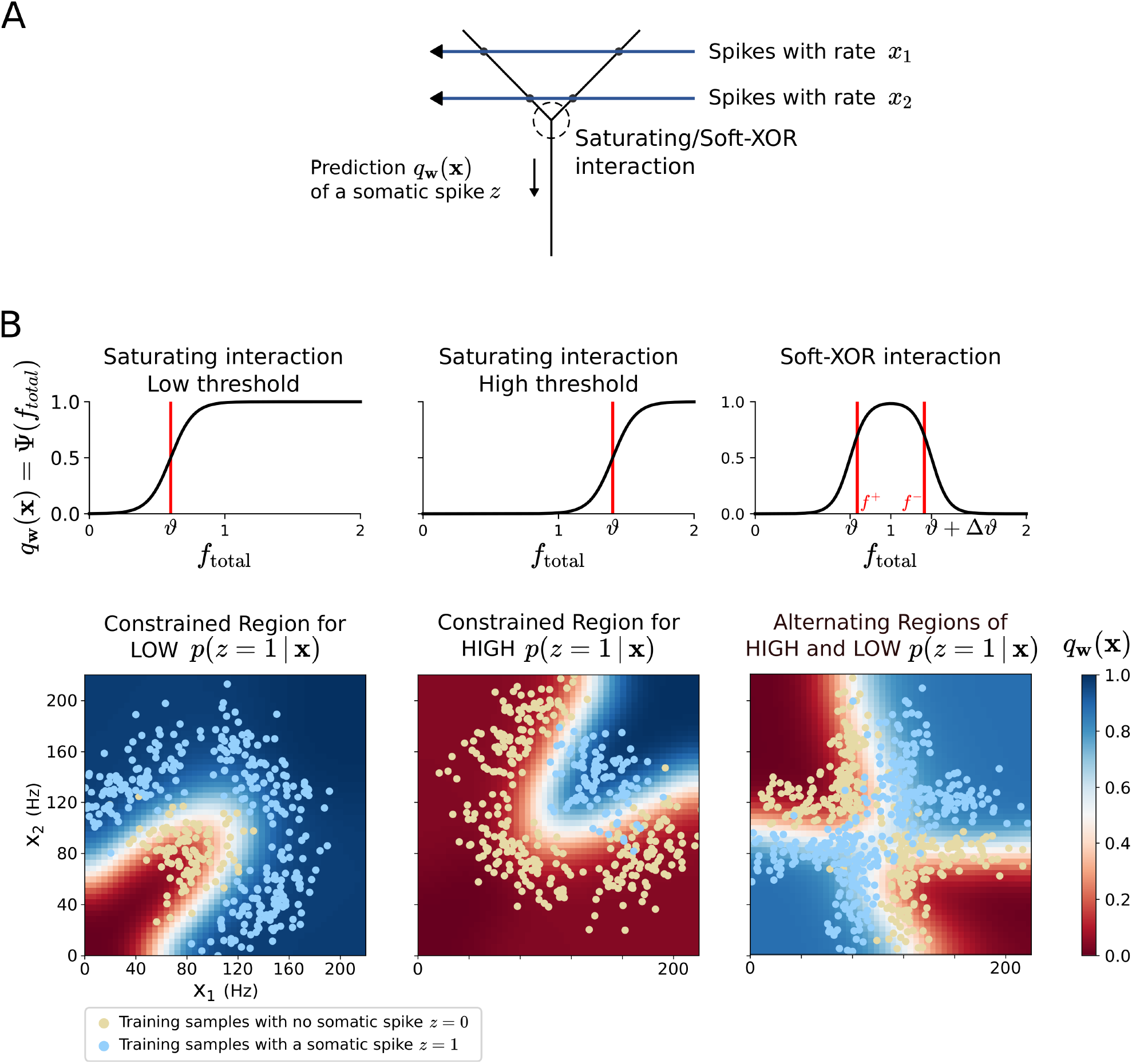
Illustration of complex probability landscapes that can be learnt in the apical dendrites of a neuron if it has more than a single branch. **A)** We consider for simplicity the case of just two branches in the dendritic tuft, but the theory also covers the case of any number of branches. We consider two different experimentally reported types of interaction between these branches: saturating and soft-XOR. **B)** The resulting probabilistic predictions Ψ(*f*_total_) of the apical tuft are shown after training on three different datasets. These were generated through mixture of Gaussians (4–8 depending on the dataset, details in Supplement). A single apical dendrite, that produces probability landscapes of the type shown in Fig. 2, cannot produce good probabilistic predictions for them. Saturating (see first two columns) and XOR-like interactions (see 3rd column) between different dendritic branches enable DLR to produce different types of probability landscapes for predicting the probability of a somatic spike. See (Table 2) for quantitative data on approximation performance.

The goal of the learning rule in this case is to get *q*_**w**_ (**x**) to predict the probability *p*_target_(*z* = 1|**x**) of somatic spikes. Hence a normative model for synaptic plasticity in these dendritic branches should minimize the distance between these two terms, formulated again as cross-entropy error.

The above choices for *f^B^* and Ψ lead to the following weight update rule for the incoming weights to a particular dendritic branch *B*:

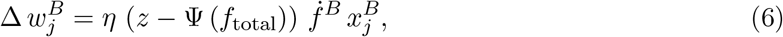

where *η* is a learning rate and 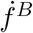 is the derivative of *f^B^* at the current value of **w**^*B*^ · **x**^*B*^. We mention two differences to the single-dendrite update: (1) the term 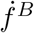 means that there is a nonlinear dependence on the local weighted input, and (2) the update depends on Ψ (*f*_total_). This quantity is not local to the dendritic branch. In order to minimize the information about Ψ (*f*_total_) at the branch, one can consider a gating of updates based on thresholds on Ψ (*f*_total_) which approximates the term (*z* – Ψ (*f*_total_)) in the update rule 6. This leads to the following updates of synaptic efficacies:

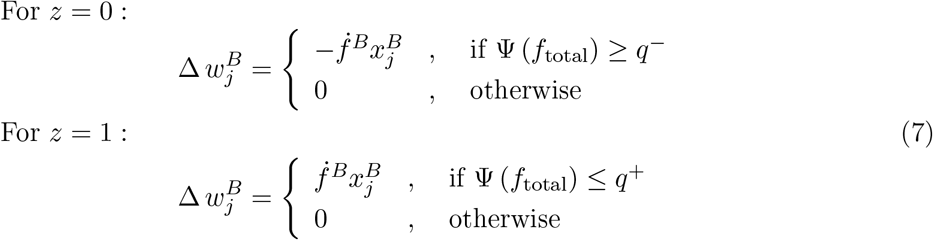

for plasticity thresholds *q*^−^ and *q*^+^. To see the correspondence with rule 6, first consider the case without somatic spike (*z* = 0). Here, the term (*z* – Ψ (*f*_total_)) in equation 6 is between −1 and 0. It becomes very small when Ψ (*f*_total_) is close to zero. This is approximated by a gating mechanism which blocks plasticity if Ψ (*f*_total_) is below some threshold *q*^−^ close to 0. When there is a somatic spike (*z* = 1), the term (*z* – Ψ (*f*_total_)) is between 0 and 1. It becomes very small when Ψ (*f*_total_) is close to one. Hence, plasticity is blocked if Ψ (*f*_total_) is above some threshold *q*^+^ close to 1 in the approximation 7.

#### 2.2.2 Soft-XOR model

This second choice for Ψ is motivated by recent findings from human L2/3 pyramidal cells. According to [Gidon et al., 2020], Ca^2+^-spikes are initiated in these cells at a threshold activation *ϑ* of dendrites, but if dendrites are even more strongly activated, these spikes are not elicited. We can model such behavior by a nonlinearity Ψ(*f*_total_) that is close to 1 in a certain activation region (*ϑ*, *ϑ* + Δ*ϑ*), see Fig. 4B (right) and Methods.

A plasticity rule for this model is derived in Methods, where the plasticity at the synapse depends on *f*_total_. Again, we can consider a coarse influence of *f*_total_ on the local plasticity, where it switches between LTP, LTD, and no update based on three thresholds *f*^−^ (low *f*_total_), *f*^mid^ (medium *f*_total_), and f^+^ (high *f*_total_), see Fig. 4B (right). This leads to the following updates of synaptic efficacies:

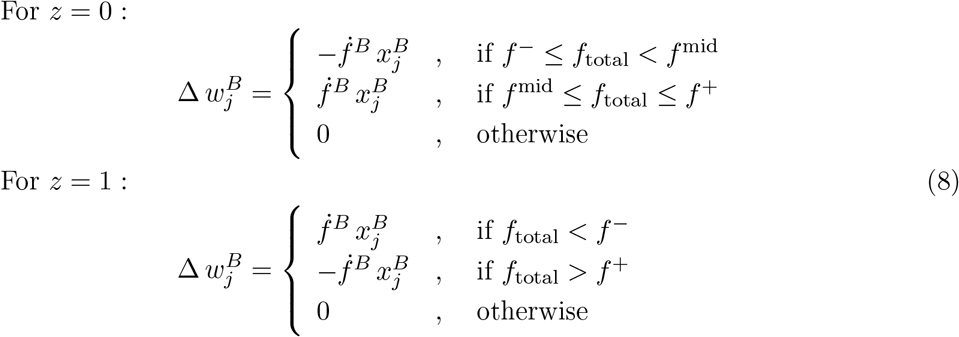

We can interpret these plasticity dynamics as follows. Consider first the case of a somatic spike *z* = 1. For low *f*_total_ (< *f*^−^), the activation has to increase in order to increase the probability of the model. Hence active synapses are potentiated. For medium *f*_total_ (*f*^−^ ≤ *f*_total_ ≤ *f*^+^), we have already a high probability for a dendritic spike. Hence, no update is necessary. For high *f*_total_ (> *f*^+^), the activation is too high and depression is needed. Consider now the case of no somatic spike *z* = 0. If the neuron is at medium *f*_total_, the probability can be decreased either by decreasing the activation (LTD) or increasing it (LTP). In the upper half of the “high probability bump” (*f*^mid^ < *f*_total_ ≤ *f*^+^), LTP is chosen, and in the lower half (*f*^−^ ≤ *f*_total_ < *f*^mid^) LTD is chosen, which shift *f*_total_ in the direction leading to a decreased probability. Outside of this range, i.e. for *f*_total_ > *f*^+^ and *f*_total_ < *f*^−^, the model already predicts a low probability for a somatic spike and thus no updates are necessary.

#### 2.2.3 Demonstration of the enhanced probabilistic prediction capabilities that emerge through DLR in the case of several apical dendrites

Similar to as shown in Fig. 2 we assume here that data points with labels *z* = 0 and *z* = 1 are generated by overlapping probability distributions. But in contrast to Fig. 2, we assume here that each of these distributions is a mixture of Gaussians, rather than single Gaussians that can be separated by an optimal linear decision boundary. Examples for the resulting clouds of data points are shown for three examples of more complex distributions in Fig. 4B.

Using a model with two dendritic branches, we consider three different kinds of interactions between these two branches – a saturating interaction with a low threshold (*ϑ* = 0.6; Fig. 4B top left), a saturating interaction with a high threshold (*ϑ* = 1.2; Fig. 4B top middle), and a soft-XOR interaction (Fig. 4B top right). One sees in Fig. 4B that each of these three types of interactions is useful for learning probabilistic predictions for three different types of distributions of data points. Furthermore one sees that training via DLR enables the dendritic branches to also provide meaningful predictions for presynaptic firing rates **x** that never occurred during training.

A detailed quantitative evaluation is provided in Methods, see Table 2 there. Here we summarize these results: Each task has regions where *p*(*z* = 1|**x**) is close to one (HIGH-*p* regions) and regions where it is close to zero (LOW-*p* regions). We tested different configurations on the following three tasks: constrained LOW-*p* region, constrained HIGH-*p* region, and alternating HIGH and LOW-*p* regions (see Fig. 4). When using two dendritic branches, a saturating interaction with low or high threshold performs well on tasks with a constrained LOW-*p* or HIGH-*p* region respectively. On the other hand, for the task with alternating LOW and HIGH-*p* regions, the Soft-XOR interaction performs much better than the saturating interaction. This demonstrates the variety of distributions that can be modeled by different configurations.

**Table 2:**
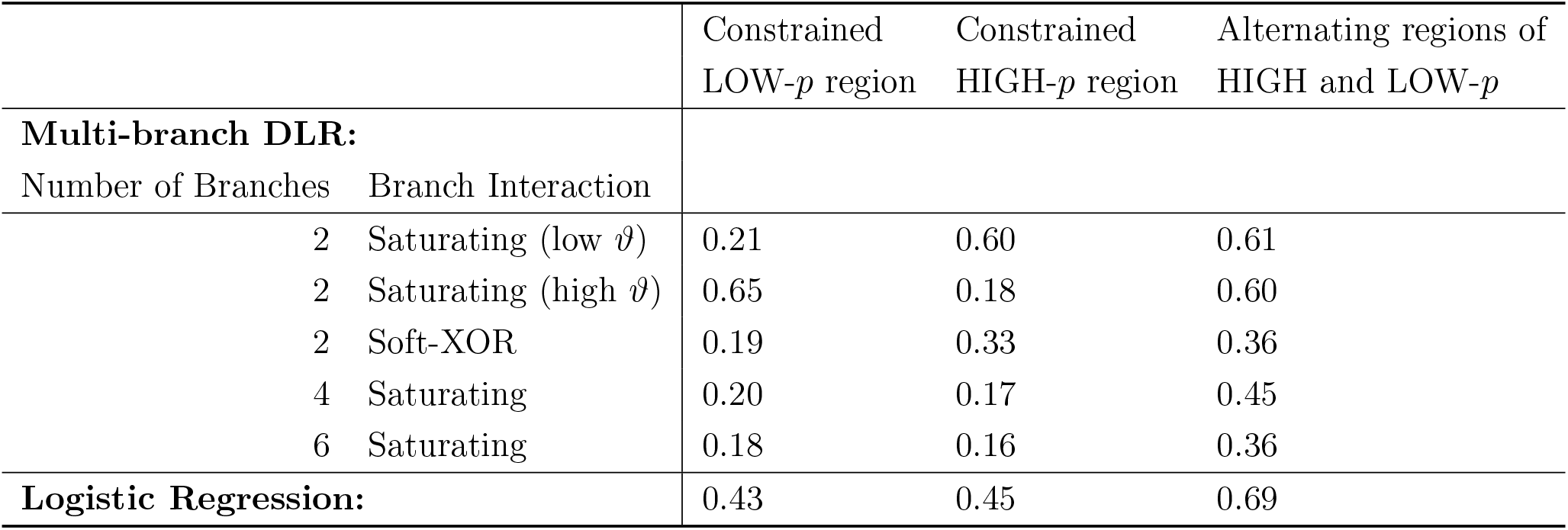
Performance of DLR for the case of several interacting apical dendrites for probabilistic predictions in cases where there are no linear optimal decision boundaries: Mean negative log likelihood (NLL) in nats (lower is better) is given for the training sets of the three different data distributions considered in Fig. 4B, and for the three types of interactions among apical dendrites that we consider, as well as for different numbers of apical dendrites. One sees that using a saturating interaction with low threshold for two branches excels at the task with a constrained LOW-*p* region, while the high threshold scheme excels at the task with a constrained HIGH-*p* region. The Soft-XOR interaction turns out to be substantially more effective for learning with two branches the probabilistic prediction for the case with alternating HIGH and LOW-*p* regions. However, with more branches the saturating interaction supports learning of probabilistic predictions via DLR for all three tasks. Each of the mean NLL values above is calculated from the best 8 out of 14 independent training runs (to avoid outliers which do not converge).

| | | Constrained<br>LOW-$p$ region | Constrained<br>HIGH-$p$ region | Alternating regions of<br>HIGH and LOW-$p$ |
| --- | --- | --- | --- | --- |
| **Multi-branch DLR:** |  |  |  |  |
| Number of Branches | Branch Interaction |  |  |  |
| 2 | Saturating (low $\vartheta$) | 0.21 | 0.60 | 0.61 |
| 2 | Saturating (high $\vartheta$) | 0.65 | 0.18 | 0.60 |
| 2 | Soft-XOR | 0.19 | 0.33 | 0.36 |
| 4 | Saturating | 0.20 | 0.17 | 0.45 |
| 6 | Saturating | 0.18 | 0.16 | 0.36 |
| **Logistic Regression:** |  | 0.43 | 0.45 | 0.69 |

We next tested whether the addition of more dendritic branches could improve the capabilities of the apical branches to predict complex distributions. To this end, we performed simulations with four and six branches using saturating interactions with intermediate threshold values (*ϑ* = 1.4 and *ϑ* = 2.1 for 4 and 6 branches respectively). We found that these configurations perform very well across all the three datasets, see Table 2. This shows that increasingly complex distributions can be modelled by a single configuration given enough branches.

## 3 Methods

### 3.1 Logistic regression

We are referring to Section 4.3.2 of [Bishop, 2006] for an introduction to logistic regression. A more informal account is given on wikipedia. It is explained there that in statistics, the (binary) logistic model is a statistical model that models the probability of one event (out of two alternatives) taking place by having the log-odds (the logarithm of the odds) for the event be a linear combination of one or more independent variables (“predictors”). In regression analysis, logistic regression is estimating the parameters of a logistic model (the coefficients in the linear combination). Formally, in binary logistic regression there is a single binary dependent *z*, while the independent variables can each be a binary variable or a continuous variable (any real value). The parameters of a logistic regression are most commonly estimated by maximum-likelihood estimation. This does not have a closed-form expression, unlike linear least squares.

Thus logistic regression provides a gold standard for predicting the events where a binary variable *z* assumes value 1, but it requires off-line computation for a batch of data. However, we show in the following subsection that the online plasticity rule DLR converges through stochastic gradient descent to the same weight vector **w**.

### 3.2 Derivation of DLR for a single apical dendrite

We denote the joint distribution of the top-down input **x** and the somatic spikes *z* as *p*_target_(**x**, *z*). Our regression formulation then seeks to provide an estimator *q*_**w**_(**x**), parametrized by **w**, where we aim to train the weights **w** so that *q*_**w**_(**x**) approximates *p*_target_(*z* = 1|**x**), that is, the probability of a somatic spike *z* given a particular top-down input **x**.

The cross entropy loss function, that measures the distance between *q*_**w**_(**x**) and the data distribution *p*_target_(*z*|**x**) is defined by

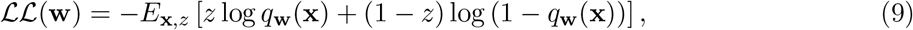

where **x**, *z* are sampled from *p*_target_(*z*, **x**).

The corresponding gradient with respect to the parameters **w** is given by

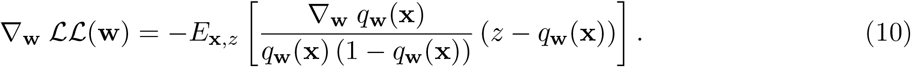

If *q*_**w**_(**x**) has the form *q*(**w** · **x**) for some nonlinearity *q*, one gets

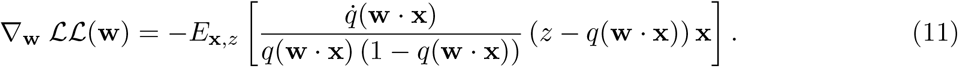

A further simplification arises if one uses a logistic sigmoid function as nonlinearity *q*:

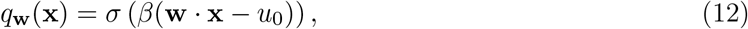

where 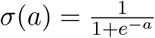 and *β* and *u*_0_ are model constants. The probability of an NMDA spike *q*_**w**_(**x**) is thus a saturated function of the input to the dendrite. Then, using the fact that *σ*′(*a*) = *σ*(*a*) (1 – *σ*(*a*)), one obtains

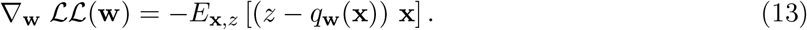

Now, *q*_**w**_(**x**) is the probability of a dendritic spike *y*. Thus we have *q*_**w**_(**x**) = *p*(*y* = 1|**x**, **w**) = *E*_*y*|**x,w**_ [*y*]. Hence we can rewrite the above rule as

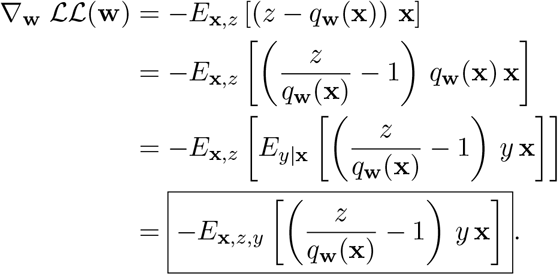

Setting the weight update as 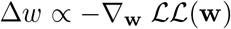 completes the derivation of the weight update for DLR described in equation 2.

This derivation proves that DLR performs stochastic descent along the gradient of the convex logistic cross entropy loss function. The convexity of this loss function guarantees stochastic convergence of the weights through DLR to the global optimum of the loss function.

### 3.3 Derivation of DLR for the case of multiple branches

Here, **x**^*B*^, **w**^*B*^ represent the weights and input to dendritic branch *B*, and **x**, **w** represent the full top-down input and weights. The corresponding cross entropy loss with *q*_**w**_(**x**) defined as in equation 3 is given by

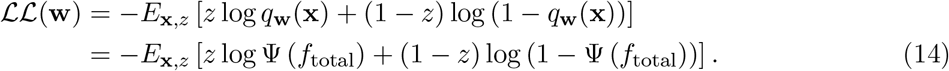

The derivative of the above loss with respect to the weights **w**^*B*^ of a branch *B* evaluates to

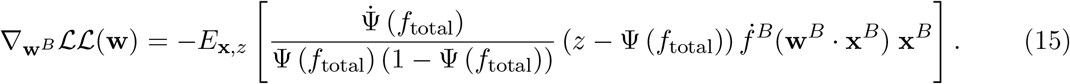

#### 3.3.1 Derivation of DLR for the saturating model

In this model, a sigmoidal shape for Ψ is chosen:

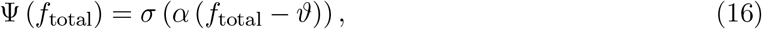

where *α* and *θ* are suitable constants. Using this in equation 15 we obtain the gradient for the saturating model:

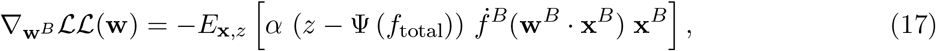

which leads to the weight update rule in equation 6.

In order to minimize the information about Ψ (*f*_total_) at the branch, one can consider a gating of updates based on thresholds on Ψ (*f*_total_) which approximates the term (*z* – Ψ (*f*_total_)) in the update rule 6. We consider the two cases *z* = 0 and *z* = 1 separately. For *z* = 0, the term (*z* – Ψ (*f*_total_)) is between −1 and 0. It becomes very small when Ψ (*f*_total_) is close to zero. A gating mechanism which blocks plasticity if Ψ (*f*_total_) is below some threshold *q*^−^ close to 0 approximates this term. For *z* = 1, the term (*z* – Ψ (*f*_total_)) is between 0 and 1. It becomes very small when Ψ (*f*_total_) is close to one. A gating mechanism which blocks plasticity if Ψ (*f*_total_) is above some threshold *q*^+^ close to 1 approximates this term. Hence, we can formulate the approximate update 7.

#### 3.3.2 Derivation of DLR for the soft-XOR model

In the soft-XOR model, Ψ is given by

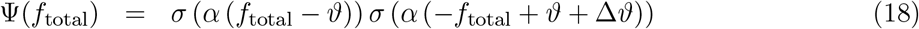

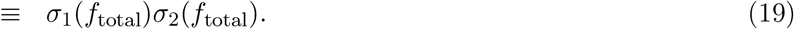

Suitable parameters *α*, *θ*, and Δ*θ* lead to an XOR-like behavior (high probability in a certain activation region), see Fig. 4B (right). Using this nonlinearity in equation 15, we obtain

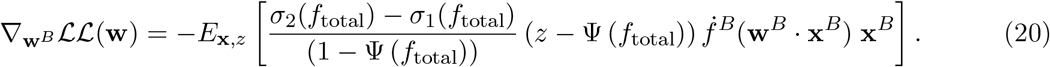

Considering a coarse influence of the *f*_total_ on the local plasticity, we can formulate an approximate rule which switches between LTP, LTD, and no update based on three thresholds *f*^−^ (close to *ϑ*), *f*^mid^ (*ϑ* + Δ*ϑ*/2), and *f*^+^ (close to *ϑ* + Δ*ϑ*), see Fig. 4B. This leads to the updates given in equation 8.

### 3.4 Simulation Details: Classification of 2D data (Fig. 2)

### 3.5 Implementation Details of DLR

For the case of a single dendritic branch, it receives top-down input spike rates **x** weighted by input weights **w**. An input vector **x** = [*x*_1_, …, *x_N_*]^*T*^ is presented to the branch as input spike trains *s*_1_(*t*), …, *s_N_*(*t*) for a fixed duration *T*_sim_. Mathematically, we model the spike trains *s_n_*(*t*) as sums of Dirac delta pulses at spike times. For each *n*, the spike times are generated by a Poisson point process of rate equal to *x_n_*. We then define the quantity

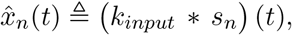

as the un-weighted input contribution through synapse *n*. where the input spikes are smoothed by the following double exponential input kernel :

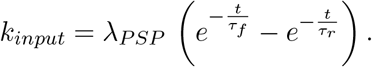

In our simulations, we used values *λ_PSP_* = 1, *τ_f_* = 2 ms, and *τ_r_* = 10 ms for the scaling- and time-constants.

It is then seen that the time average of 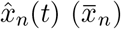, taken over the interval of presentation *T*_sim_, is proportional to the input spike rate **x**. Thus, for the weight update rule in 2, we use the above time average 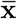 as a proxy for **x** to calculate *q*_**w**_(**x**). This is because 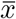 is a quantity that is local to the incoming dendritic synapse and can thus be used in local synaptic weight updates.

For the case of the single branch SLR, the implemented weight update is thus given by

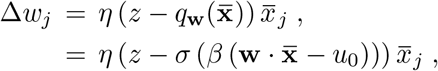

where we make use of the definition of *q*_**w**_(**x**) from equation 1.

Similarly, in the case of multi-branch update rules, for each branch *B*, we calculate using a spike trace 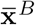 as an estimate proportional to **x**^*B*^ and use 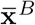 when implementing the weight update rules corresponding to the saturating and soft-XOR dynamics in equations 7, 8.

#### 3.5.1 The target distribution *p*_target_(**x**, *z*)

We first describe the distribution of the training data. The training examples for this task consist of points generated from two 2D Gaussian clusters with means and covariances ***μ**_k_*, **Σ**_*k*_ for *k* ∈ {0, 1}. All points generated from 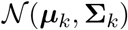 have the target value *z* = *k*. The prior probabilities for the classes 0 and 1 were chosen to be equal (0.5). Thus to generate a training sample (**x**_*i*_, *z_i_*), we first pick the Gaussian component *k* ∈ {0, 1} with equal probability i.e. *k* ~ bernoulli(*p* = 0.5). We then pick 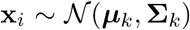 and assign the target *z_i_* = *k*. This gives the following conditional probability distribution *p*_target_(*z* = 1|**x**)

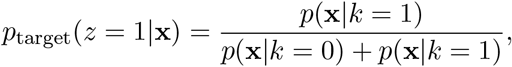

where *p*(**x**|*k*) is the Gaussian probability density function with mean ***μ**_k_* and covariance **Σ**_*k*_. It is the aim of the algorithm to learn this conditional distribution in the quantity *q*_**w**_(**x**) in the dendrite (equation 1).

The covariance matrix **Σ**_*k*_ can be decomposed as 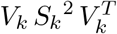. Here *V_k_* is an orthogonal matrix where the columns are the eigenvectors of **Σ**_*k*_ and give the directions along which the 2D Gaussian is aligned. 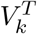 denotes the transpose of the matrix. *S_k_* is a diagonal matrix with each diagonal entry representing the standard deviation of the Gaussian along the corresponding eigenvector. The parameters used for this simulation are:

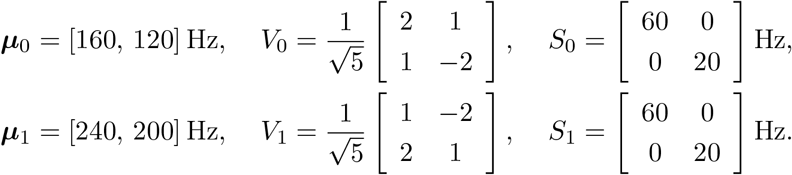

#### 3.5.2 Input encoding and training

The neuron received the input on a single dendritic branch, with the DLR being applied to the incoming weights to that branch. Each data point **x**_*i*_, is padded with an additional component of constant rate of 40Hz (in order to model intercept fitting). Each component of this data point is then given as input to the network encoded as a Poisson spike train with mean rate equal to the value of the component. This spike train is provided to the neuron for a duration of *T*_sim_ = 400*ms*. Targets are fed to the network by inducing or preventing somatic spikes corresponding to *z* = 1 and *z* = 0 respectively. Fig. 1B illustrates this neuron and learning setup.

When training the input weights to the neuron using DLR, the initial weights for each run are initialized to positive values drawn from a normal distribution with mean 9.0 and standard deviation 4.5. The weights are clipped at zero so that they stay positive in order to obey Dale’s law.

The standard perceptron classifier trains a set of weights **w** so that for any data point (**x**, *z*), we should have **w** · **x** > 0 if *z* = 1 and **w** · **x** < 0 if *z* = 0. For each iteration of updating the weights **w**, we randomly sample a single data point (**x**, *z*) from the target distribution *p*_target_, and apply the following standard perceptron update to weights **w**. Note that we also add here the same additional component of 40Hz to **x** to enable the perceptron to train the intercept of the separating line. The perceptron update rule is provided below

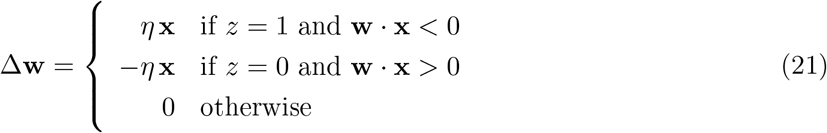

The learning rates *η* for both the DLR and perceptron learning (used in equations 2, and 21 respectively) decay harmonically from an initial value of *η*_0_ to a final value of *η_final_* over the course of the 30000 iterations that we run them for in Fig. 2. For the DLR we use *η*_0_ = 1.8 (See Table 3), and for the perceptron learning we use *η*_0_ = 0.01, where the lower learning rate is used to aid any possible convergence of the perceptron. In both cases *η_final_* = *η*_0_/6. Thus we see that even with a lower learning rate and an idealized perceptron learning, the perceptron rule does not converge for this case where the data is not linearly separable.

**Table 3:** Parameters related to DLR that are used for each experiment. *η*_0_, *η*_final_ are the initial and final learning rate used. *β* and *u*_0_ are the scale and threshold for the sigmoid function used to calculate *q*_**w**_(**x**) and *f*^*B*^ in equations 1 and 4 respectively.

| Parameter | Single-branch DLR |  | Multi-branch DLR |  |
| --- | --- | --- | --- | --- |
|  | 2-D regression | Predicting input from moving object | Saturating interaction | Soft-XOR interaction |
| $\eta_0$ | 1.8 | 40.0 | 4.0 | 0.3 |
| $\eta_{final}$ | 0.3 | 6.67 | 4.0 | 0.3 |
| $\beta$ | 0.5 | 0.5 | 0.1 | 1.0 |
| $u_0$ | 20.0 | 15.0 | 7.0 | 7.0 |
| $\lambda_{PSP}$ | 1.0 | 1.0 | 1.0 | 1.0 |
| $\tau_r$ | 2.0 ms | 2.0 ms | 2.0 ms | 2.0 ms |
| $\tau_f$ | 10.0 ms | 10.0 ms | 10.0 ms | 10.0 ms |
| $T_{sim}$ | 400.0 ms | 200.0 ms | 400.0 ms | 400.0 ms |

Note that incoming weights are initialized to random positive values and the sign of the weight is fixed in accordance to Dale’s law

### 3.6 Simulation Details: Predicting the future sensory input from a moving object (Fig. 3)

#### 3.6.1 Input distribution

We consider here the following task. Given estimates of the position and velocity of a moving object at time 0, predict the sensory input expected from the object at its new position after a fixed delay of *T*.

In our simulation, the moving object is a 2D circle which moves within a 30 × 30 grid and has a diameter of 5 Grid Units (GU). More specifically, we have a 30 × 30 grid of pyramidal neurons, each of which correspond to one position in the aforementioned grid on which the object moves. The soma of each neuron spikes (i.e. *z* = 1) iff the object covers the corresponding position on the grid.

The information that forms the dendritic input is given by the four-tuple (*r_x_, r_y_, v, θ*) consisting of x and y coordinates *r_x_*, *r_y_* of the initial position, the speed *v*, and the direction angle *θ*. The initial position (*r_x_*, *r_y_*) can take any value such that both *r_x_*, *r_y_* take values in the range of *S_r_* = [3, 27] GU. The speed v is defined in terms of the distance (in GU) that the moving object covers in the fixed delay *T* between the top-down and the bottom up inputs. This distance can take any value in the range *S_v_* = [4, 20] GU. The direction is specified by an angle *θ* to the horizontal which can take any value in the range [0, 2*π*).

In this setup each data point is generated as follows. We sample a four-tuple (*r_x_, r_y_, v_real_, θ_real_*) picking each parameter uniformly from the above ranges. This four-tuple represents the initial position and real velocity of the object at the initial time point *t* = 0. In order to model the noise introduced by the higher layers, we only feed a noisy perturbation of the real velocity which is calculated as follows

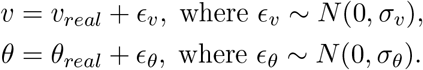

The input spike rates **x** is the population-coded represenation of (*r_x_, r_y_, v, θ*), where the population coding is described below. For each pyramidal neuron, the presence of a somatic spike *z* is determined by the image of the object on the grid at time *t* = *T*, which is calculated using (*r_x_, r_y_, v_real_, θ_real_*). A pyramidal neuron will have a somatic spike (*z* = 1) iff in this image, the object covers the corresponding grid position. With this, the incoming weights to each pyramidal neuron are trained using DLR with the target being the corresponding pixel value in this image.

Given the perturbed top-down estimate of initial position and velocity at time 0 as input, the true initial velocity of the object, and consequently the final position of the object at time T is uncertain. The calculation of the corresponding target distribution *p*_target_(*z* = 1|**x**) is described below in Section 3.6.3.

#### 3.6.2 Input encoding and parameters

The top-down inputs specifying the initial position and velocity of the object are encoded into neuron populations as follows. We consider the sets *M_r_*, *M_v_* and *M_θ_*, which consist of *N_r_*, *N_v_*, and *N_θ_* equally spaced values in the ranges *S_r_*, *S_v_*, and *S_θ_* respectively. Then, corresponding to each four-tuple (*μ_x_*, *μ_y_*, *μ_v_*, *μ_θ_*) ∈ *M_r_* × *M_r_* × *M_v_* × *M_θ_*, we have a single input neuron that fires according to a Gaussian tuning curve centered at (*μ_x_, μ_y_, μ_v_, μ_θ_*). The neuron encodes the input (*r_x_, r_y_, v, θ*) by firing at a rate *ρ* given by

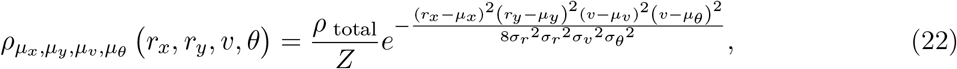

where *Z* is a normalization factor calculated so that the total spike rate of all the input neurons is *ρ* _total_ = 250 Hz. Here the standard deviation of the response curve corresponding to position *σ_r_*, velocity *σ_v_*, and direction *σ_θ_* are constant across all neurons. **x** is the set of all such input neurons.

For the simulation whose results are shown in Fig. 3, we use *N_r_* = 10, *N_v_* = 10, *N_θ_* = 24, *σ_r_* = 2.67, *σ_v_* = 1.78, *σ_θ_* = 0.2618 = 15.0°. This leads to **x** having a total dimension of *N_r_*^2^*N_v_N_θ_* = 24000. The standard deviations of the Gaussian noise perturbations applied to the speed and direction components of the top-down input are equal to the width of the respective receptive fields *σ_v_, σ_θ_*. In order to train the network we make use of 168000 data points in the training dataset.

All the weights are initialized to a constant positive value of 9.0 and are clipped at zero during training so that they stay non-negative in accordance with Dale’s law.

#### 3.6.3 Calculating actual probability of a neuron’s receptive field being covered by the object at time *T*

The imprecision in the estimate of the object velocity, modeled by the perturbed input speed and direction, leads to the final position of the object being uncertain. Each pyramidal neuron in the 30 × 30 grid receives as input to the apical dendrites the four-tuple 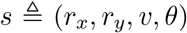. Here (*r_x_, r_y_*) is the initial position of the object and (*v, θ*) are the noisy estimates of speed and direction of the velocity used by the pyramidal neuron. We denote the true velocity of the object by (*v_real_, θ_real_*), and define the corresponding four-tuple 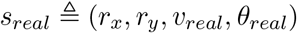.

Consider a single neuron in the grid. For this neuron, the soma spikes (*z* = 1) if and only if the object covers the receptive field of this neuron at time *T*, and *z* = 0 otherwise. Given the noise in the estimate of velocity, we thus wish to calculate *p*_target_(*z* = 1|**x**), which is the conditional distribution that is estimated by the neuron via *q*_**w**_ (**x**)

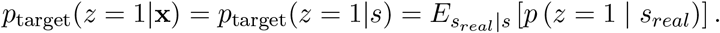

Here we use the fact that the final position of the object is fully determined given the true velocity and the initial position *s_real_*.

To calculate *p*(*s_real_* |*s*) we then use Bayes’ theorem:

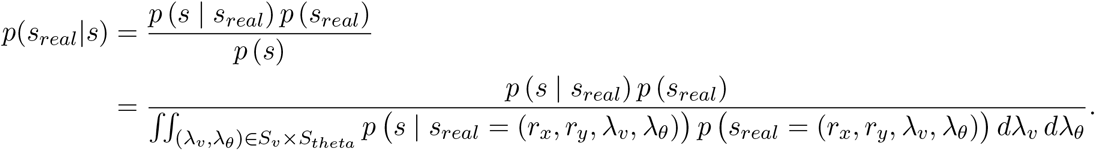

Since *s_real_* is sampled uniformly, *p*(*s_real_*) is a constant, hence it simplifies to

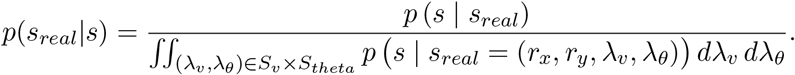

The above can be computed using the fact that *p*(*s*|*s_real_*) is the probability density function of a normal distribution.

#### 3.6.4 Measurement of performance

Here we provide measurements to demonstrate that the network reliably predicts the probability that a neuron receives input from the moving object at time *T*. We term the position of the center of the moving object as predicted by the noisy estimate of the velocity as the *estimated final position*. We then consider three concentric slices of the neuron grid centered at the estimated final position, with diameters 0 – 5 GU, 5 – 9 GU, and 9 – 13 GU. For the pyramidal neurons within a slice, we compare the mean predicted probability *q*_**w**_(**x**) and the mean actual probability, of their receptive fields being covered by the object at time *T*.

1. Slice of diameter 0 – 5 GU: mean *q*_**w**_(**x**) = 0.3546, mean actual probability = 0.4108
2. Slice of diameter 5 – 9 GU: mean *q*_**w**_(**x**) = 0.1806, mean actual probability = 0.1668
3. Slice of diameter 9 – 13 GU: mean *q*_**w**_(**x**) = 0.0638, mean actual probability = 0.0435

From this we can clearly see that the mean predicted probability *q*_**w**_ (**x**) drops away from the estimated final position, as well as that the theoretically calculated probabilities are followed closely by the predicted probabilities. The examples in Fig. 3 and the results above are calculated from combinations of initial position and velocity that were not seen during training.

### 3.7 Simulation Details: Predicting complex distributions with multiple dendritic branches (Fig. 4)

#### 3.7.1 Target distribution *p*_target_(x, *z*)

In this experiment, the 2D data is generated from multiple gaussian clusters. Each gaussian cluster *k* has a mean ***μ**_k_*, covariance **Σ**_*k*_, and a corresponding value of the somatic spike *ζ_k_* ∈ {0, 1}. To generate a data point, we first pick a cluster *k* ∈ 1, …, *K* at random, sample **x** from it, and assign the somatic spike *z* = *ζ_k_* corresponding to the cluster. The resultant joint target distribution is

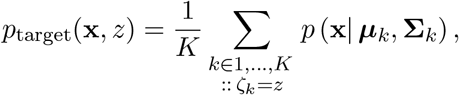

where *p*(**x**| ***μ**_k_*, **Σ**_*k*_) is the gaussian probability density function with mean ***μ***_*k*_ and covariance **Σ**_*k*_.

The parameters and the somatic spiking corresponding to the different clusters are chosen so as to create three distributions which have respectively a constrained LOW-*p* region, constrained HIGH-*p* region, and alternating HIGH and LOW-*p* regions (see Fig. 4). Here LOW and HIGH-*p* refer to regions of LOW and HIGH *p*_target_(*z* = 1|**x**) respectively. The precise parameters of the component gaussian distributions are provided in the Supplement.

#### 3.7.2 Input encoding and training parameters

In order to model the above distributions, we consider a neuron with *N_B_* (two or more) dendritic branches each with their respective incoming synapses with weights **w**^1^, …,**w**^*N_B_*^. All of these branches receive the same 2D input, i.e., **x**^*B*^ = [**x**; 1] = [*x*_1_, *x*_2_, 1] for *B* = 1, …, *N_B_*. This input is provided as in the 2D regression task for a single branch, i.e., as a poisson spike sequence with spike rates given by **x**^*B*^. The additional input component with a value of 20 Hz is given in order to model the training of the branch baseline activation.

The various configurations in Table 2 along with corresponding parameters are below.

1. 2 Branches, saturating interaction, low threshold *ϑ* = 0.6: This implements a soft-OR operation between the two branches.
2. 2 Branches, saturating interaction, high threshold *ϑ* = 1.2: This implements a soft-AND operation between the two branches.
3. 2 Branches, soft-XOR interaction, *ϑ* = 0.5, Δ*ϑ* = 1.0. This implements a soft-XOR operation between the two branches.
4. 4 Branches, saturating interaction, *ϑ* = 1.4.
5. 6 Branches, saturating interaction, *ϑ* = 2.1.

For all the saturating interactions, we use a scale factor *α* = 12. The parameters *q*^+^, *q*^−^ used in the learning rule in 7 are set to *q*^+^ = 0.9 and *q*^−^ = 0.1. For the Soft-XOR interaction, we use *α* = 8.0 and set *f*^−^ = *ϑ* = 0.5, *f*^+^ = *ϑ* + Δ*ϑ* = 1.5 and *f*^mid^ = *ϑ* + Δ*ϑ*/2 = 1.0.

The initial weights for each branch *B* are initialized in the following manner. We pick a point 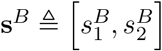, where both 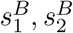 are randomly picked uniformly from the interval [110, 130] Hz. We then pick a random direction vector **h**^B^ sampled uniformly from a unit circle. Using this pair of **s**^*B*^, **h**^*B*^, we set the initial weights 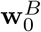 so that the separating plane is a plane that passes through the point **s**^*B*^, with the direction of the plane determined by the normal vector **h**^*B*^. The weights in this case are unconstrained in sign to allow the model to fit the complex distributions in the case of the low dimensional inputs.

### 3.8 Common Parameters for DLR

Some of the common parameters and the values for the DLR rule for the three experiments are given in Table 3.

## 4 Discussion

We have provided a link between the probabilistic theory of [Lee and Mumford, 2003] for computations in hierarchical cortical networks and synaptic plasticity in apical dendrties of pyramidal cells on layers 2/3 and 5 of generic cortical microcircuits. It has recently been proposed [Shin et al., 2021] that these pyramidal cells become selective for specific bottom-up and top-down input features in two separate learning phases: During the first learning phase these neurons become selective to specific features of sensory inputs, or more generally to specific features of the bottom-up input stream which reaches them primarily through synapses on basal dendrites. Synaptic plasticity on apical dendrites enables these neurons during the second learning phase to integrate their bottom-up processing into concrete contexts and goals. We have presented here a learning theory for this second learning phase. More precisely, we have shown that the theory of [Lee and Mumford, 2003], which postulates a concrete computational goal as outcome of this second learning phase -top-down generation of probabilistic priors for bottom-up information streams, is in many aspects consistent with known experimental data on synaptic plasticity in apical dendrites. More precisely, we have presented a normative model for these plasticity processes based on the hypothesis that their functional goal is self-supervised learning of probabilistic priors from the statistics of coincidences and non-coincidences of particular configurations of firing patterns in the top-down and bottom-up input streams that converge on a neuron. This normative model builds on a key result from statistical learning theory [Bishop, 2006]: That logistic regression defines the optimal weight vector for producing such probabilistic prediction through a weighted sum of top-down inputs **x**, provided a sigmoid function is used to map the weighted sum into probabilities, i.e., values between 0 and 1. We have identified a concrete online synaptic plasticity rule, called Dendritic Logistic Regression (DLR), that approximates this gold standard. This plasticity rule is consistent with a number of known details of plasticity rules in apical dendrites, in particular with the prominent role of NMDA spikes for this plasticity [Gambino et al., 2014, Stein et al., 2021]. However many details of synaptic plasticity in apical dendrites remain unknown, especially with regard to LTD.

Understanding learning of such priors, i.e., probabilities, requires to step away from the more familiar set-up of learning in a deterministic setting, where the predicted “labels” are provided by a generally consistent supervisor, rather than through a stochastic process. Although one is tempted to think also about synaptic plasticity in apical dendrites in terms of well-known learning rules such as the perceptron learning rule, a closer look (see also Fig. 2 and S1) shows that such learning rule is not suitable for learning the probability of a stochastically occuring event (such as a somatic spike). The key point is that such somatic spike will sometimes occur for a given top-down input **x** and sometimes not. We have demonstrated in Fig. 2B that DLR is well-suited for coping with this challenge: It can even learn for data points that are generated by two overlapping Gaussians the probability that a data point was generated by a specific one of them. In fact DLR can learn this probability as well as logistic regression. In Fig. 3 we have scaled up self-supervised learning through DLR to a 2D sheet of pyramidal cells, and have shown that it enables them to learn a close to optimal 2D probability landscape for estimating the future position of a moving object.

A long-standing open question is whether dendrites offer to neurons a substantial functional advantage, in particular, a conceptual advantage that goes beyond integrating computational capabilities into single neurons for which otherwise a larger neural network would be needed [Larkum, 2022]. Our results provide an interesting new answer to this question: Dendritic arborization enables neurons to learn internal models (probability landscapes) that are substantially more complex, and therefore can provide substantially better fits to target distributions that are generated by their environment, see Fig. 4. We are not aware of a biologically plausible method that would enable even substantially larger networks of point neurons to achieve that, because it requires an integration of several non-binary signals that are readily available within a single neuron, but which are apparently not transmitted between neurons. For the sake of better visualization we have demonstrated in Fig. 4 this qualitative jump in the complexity of probability landscapes that can be learnt by neurons with several apical dendrites only for the case where the top-down input **x** is 2-dimensional. But the underlying theory that we have presented holds for any dimension of **x**. Obviously, a further qualitative jump in the complexity of internal probabilistic models that can be learnt arises when one applies DLR to neurons that receive 100- or 1000 dimensional top-down inputs **x**.

There exists a staggering diversity of subtypes of pyramidal cells, especially in the human cortex [Berg et al., 2021]. In addition there exists substantial diversity among subtypes of NMDA receptors [Stein et al., 2021] and of NMDA spikes [Stuyt et al., 2021, Larkum et al., 2022]. Hence it would be quite optimistic to expect that all resulting plasticity processes in apical dendrites of pyramidal cells can be described by a simple formula, such as DLR. However, a comparison of postulates of DLR with experimental data on synaptic plasticity processes in a concrete apical dendrite will provide definite insight into the question whether this plasticity process is likely to have the functional goal to learn the probability of a somatic spike. Since DLR is approximately optimal for learning this probability, significant deviations from DLR suggest that this learning process has a different functional goal.

## Acknowledgments

This work was supported by the European Union’s Horizon 2020 Framework Programme for Research and Innovation under the Specific Grant Agreements No. 785907 (Human Brain Project), the project No. 899265 (ADOPD), and by a grant from Intel.

