## Supplemental Information for "Self-supervised learning of probabilistic prediction through synaptic plasticity in apical dendrites: A normative model"

### Supplement to Self-supervised learning of probabilistic predictions of bottom-up inputs to pyramidal cells through synaptic plasticity in apical dendrites: A normative model

Arjun Rao[*,1], Robert Legenstein[*,1], Anand Subramoney[1,2], and Wolfgang Maass[1,+]

[*]First authors

[1]Institute for Theoretical Computer Science, Graz University of Technology, 8010, Austria
[2]Institute for Neural Computation, Ruhr University Bochum, Germany

June 23, 2022

#### S1 Learning a constant probability $p(z = 1|\mathbf{x})$

In this supplementary section we demonstrate that DLR can also learn probability landscapes that do not interpolate between values 0 and 1, and rather provide a flat prior. In this example we generate training data of input firing rates $\mathbf{x}$ and somatic spiking $z$ from the distribution $p_{\text{target}}(\mathbf{x}, z)$ that is defined below.

Each component of $\mathbf{x}$ is sampled uniformly from $[0, 240]$ Hz, and for any $\mathbf{x}$, the probability of a somatic spike is $p_{\text{HIGH}} = 0.3$. Mathematically we have

$$\mathbf{x} \triangleq [x_1, x_2] \text{ where } x_1, x_2 \sim \mathcal{U}\left([0, 240] \text{ Hz}\right) \tag{1}$$

$$p_{\text{target}}(z = 1|\mathbf{x}) \triangleq p_{\text{HIGH}} = 0.3 \text{ for all } \mathbf{x}. \tag{2}$$

For each data point $\mathbf{x}_i$, we provide an additional input of constant rate 40Hz (in order to model intercept fitting). The target distribution is thus not related to a classification problem and no classifying boundary would be any better than another. However DLR is able to appropriately predict $p_{\text{target}}(z = 1|\mathbf{x})$ via the value of $q_{\mathbf{w}}(\mathbf{x})$ as shown in Fig. S1.

#### S2 Details: Demonstration of the enhanced probabilistic prediction capabilities that emerge through DLR in the case of several apical dendrites

##### S2.1 Parameters of the target distributions

In this experiment, the 2D data is generated from a mixture of Gaussians, with two gaussian clusters, one for each value of z.. Each Gaussian $k$ has a mean $\mu_k$, covariance $\Sigma_k$, and a corresponding value

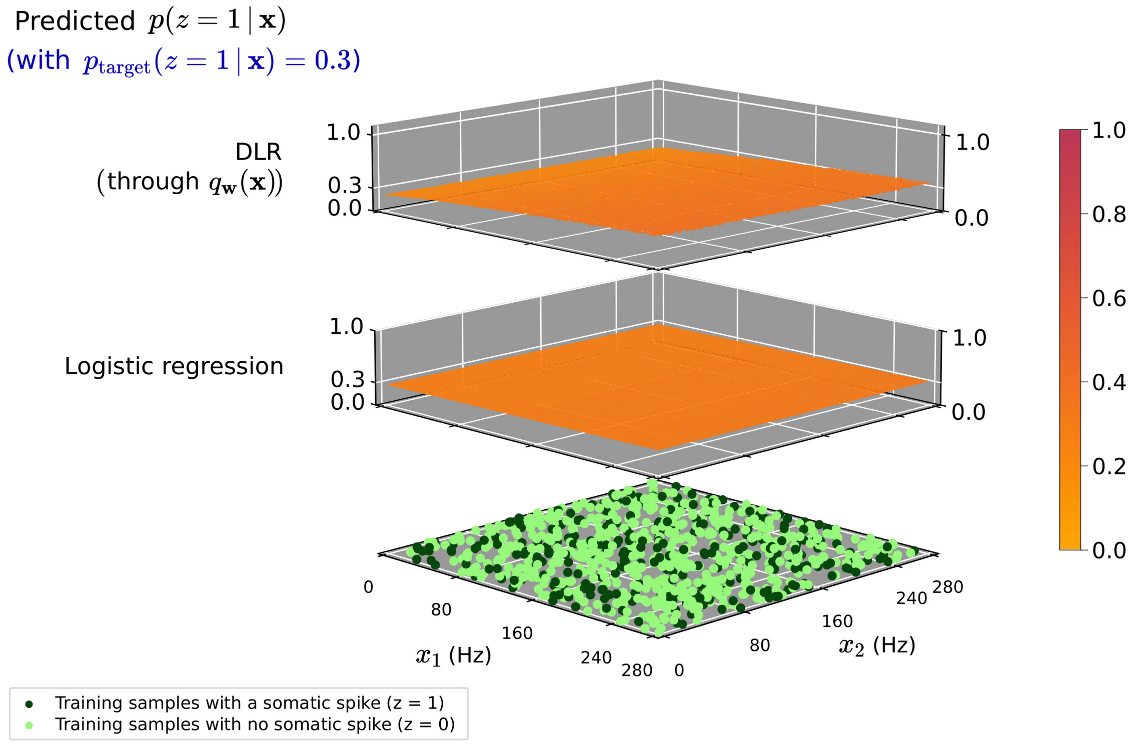


**Figure S1: Illustration of the ability of DLR to estimate probability distributions.** Shown in the bottom plane is $\mathbf{x}$ sampled from a uniform distribution, where the coordinate of each point is the input firing rate of a top-down neuron into the dendritic branch. For each point $\mathbf{x}$, the corresponding probability of a somatic spike is $p_{\text{target}}(z = 1|\mathbf{x}) = p_{\text{HIGH}} = 0.3$. The top plane shows the value of $q_{\mathbf{w}}(\mathbf{x})$ with the weights trained using DLR. We see that $q_{\mathbf{w}}(\mathbf{x})$ approximates $p_{\text{HIGH}}$ demonstrating that DLR learns probability distributions as opposed to purely a separating boundary. The corresponding predicted probability via theoretically optimal logistic regression is shown in the middle plane.

$\zeta_k$ of $z$. To generate a data point, we first pick a cluster $k \in \{1, \ldots, K\}$ at random, sample $\mathbf{x}$ from it, and assign the value $z = \zeta_k$ to $z$. The resultant joint target distribution is

$$p_{\text{target}}(\mathbf{x}, z) = \frac{1}{K} \sum_{k \in \{1,\ldots,K \mid \zeta_k = z\}} p\left(\mathbf{x} \mid \mu_k, \Sigma_k\right),$$

where $p\left(\mathbf{x} \mid \mu_k, \Sigma_k\right)$ is the Gaussian probability density function with mean $\mu_k$ and covariance $\Sigma_k$.

The shape of the Gaussian distribution for any cluster $k$ can be specified via the 2D vector $\sigma_k$ and the $2 \times 2$ matrix $V_k$. The columns of $V_k$ are unit vectors perpendicular to each other that specify the directions along which the 2D Gaussian is oriented, and $\sigma_k$ specifies the standard deviation along the respective directions. The covariance matrix $\mathbf{\Sigma}_k$ can be reconstructed as

$$\mathbf{\Sigma}_k = V_k \begin{bmatrix} \sigma_{k,1}^2 & 0 \\ 0 & \sigma_{k,2}^2 \end{bmatrix} V_k^T.$$

The parameters and the somatic spiking corresponding to the different clusters are chosen so as to create three distributions which have respectively a constrained LOW-$p$ region, constrained HIGH-$p$ region, and alternating HIGH and LOW-$p$ regions (see Fig. 4). Here LOW and HIGH-$p$

| $k$ | Somatic activity ($\zeta_k$) | $\boldsymbol{\mu}_k$ | $V_k$ | $\sigma_k$ |
|---|---|---|---|---|
| 0 | 0 | $[80, 80]$ Hz | $\frac{1}{\sqrt{2}}\begin{bmatrix} 1 & -1 \\ 1 & 1 \end{bmatrix}$ | $[28, 14]$ Hz |
| 1 | 0 | $[180, 80]$ Hz | $\frac{1}{\sqrt{2}}\begin{bmatrix} 1 & -1 \\ 1 & 1 \end{bmatrix}$ | $[14, 28]$ Hz |
| 2 | 0 | $[80, 180]$ Hz | $\frac{1}{\sqrt{2}}\begin{bmatrix} 1 & -1 \\ 1 & 1 \end{bmatrix}$ | $[14, 28]$ Hz |
| 3 | 1 | $[140, 140]$ Hz | $\frac{1}{\sqrt{2}}\begin{bmatrix} 1 & -1 \\ 1 & 1 \end{bmatrix}$ | $[20, 20]$ Hz |

**Table S1: Parameters for the distribution with a constrained HIGH-$p$ region**. Only one of the four clusters generates points with $Z = 1$ (i.e. for which $\zeta_k = 1$).

refer to regions of LOW and HIGH $p_{\text{target}}(z = 1|\mathbf{x})$ respectively. The parameters for these three distributions, i.e. $\boldsymbol{\mu}_k$, $V_k$, $\sigma_k$, $\zeta_k$ are specified in Tables S1–S3.

| $k$ | Somatic activity ($\zeta_k$) | $\boldsymbol{\mu}_k$ | $V_k$ | $\sigma_k$ |
|---|---|---|---|---|
| 0 | 0 | $[80, 80]$ Hz | $\frac{1}{\sqrt{2}}\begin{bmatrix} -1 & 1 \\ 1 & 1 \end{bmatrix}$ | $[28, 14]$ Hz |
| 1 | 1 | $[140, 40]$ Hz | $\frac{1}{\sqrt{2}}\begin{bmatrix} -1 & 1 \\ 1 & 1 \end{bmatrix}$ | $[14, 28]$ Hz |
| 2 | 1 | $[40, 140]$ Hz | $\frac{1}{\sqrt{2}}\begin{bmatrix} -1 & 1 \\ 1 & 1 \end{bmatrix}$ | $[14, 28]$ Hz |
| 3 | 1 | $[140, 140]$ Hz | $\frac{1}{\sqrt{2}}\begin{bmatrix} -1 & 1 \\ 1 & 1 \end{bmatrix}$ | $[28, 14]$ Hz |

**Table S2: Parameters for the distribution with a constrained HIGH-$p$ region**. Only one of the four clusters generates points with $Z = 0$ (i.e. for which $\zeta_k = 0$).

| $k$ | Somatic activity ($\zeta_k$) | $\boldsymbol{\mu}_k$ | $V_k$ | $\sigma_k$ |
|---|---|---|---|---|
| 0 | 1 | $[80, 50]$ Hz | $\begin{bmatrix} 1 & 0 \\ 0 & 1 \end{bmatrix}$ | $[10, 30]$ Hz |
| 1 | 1 | $[50, 80]$ Hz | $\begin{bmatrix} 1 & 0 \\ 0 & 1 \end{bmatrix}$ | $[30, 10]$ Hz |
| 2 | 0 | $[50, 120]$ Hz | $\begin{bmatrix} 1 & 0 \\ 0 & 1 \end{bmatrix}$ | $[30, 10]$ Hz |
| 3 | 0 | $[80, 150]$ Hz | $\begin{bmatrix} 1 & 0 \\ 0 & 1 \end{bmatrix}$ | $[10, 30]$ Hz |
| 4 | 1 | $[120, 150]$ Hz | $\begin{bmatrix} 1 & 0 \\ 0 & 1 \end{bmatrix}$ | $[10, 30]$ Hz |
| 5 | 1 | $[150, 120]$ Hz | $\begin{bmatrix} 1 & 0 \\ 0 & 1 \end{bmatrix}$ | $[30, 10]$ Hz |
| 6 | 0 | $[150, 80]$ Hz | $\begin{bmatrix} 1 & 0 \\ 0 & 1 \end{bmatrix}$ | $[30, 10]$ Hz |
| 7 | 0 | $[120, 50]$ Hz | $\begin{bmatrix} 1 & 0 \\ 0 & 1 \end{bmatrix}$ | $[10, 30]$ Hz |

**Table S3: Parameters for the distribution with alternating HIGH-$p$ and LOW-$p$ region**. Here clusters have alternating values of $\zeta_k$.
